# Open access policies of leading medical journals: a cross-sectional study

**DOI:** 10.1101/250613

**Authors:** Tim S Ellison, Tim Koder, Laura Schmidt, Amy Williams, Christopher C Winchester

## Abstract

**Objectives:** Academic and not-for-profit research funders are increasingly requiring that the research they fund must be published open access, with some insisting on publishing with a Creative Commons Attribution (CC BY) licence to allow the broadest possible use. We set out to clarify the open access variants provided by leading medical journals for research in general and industry-funded research in particular, and record the availability of the CC BY licence for commercially funded research.

**Methods:** We identified medical journals with a 2015 impact factor of at least 15.0 on 24 May 2017, then excluded from the analysis journals that only publish review articles. Between 29 June 2017 and 26 July 2017, we collected information about each journal’s open access policies from their websites and/or by email contact. We contacted the journals by email again between 6 December 2017 and 2 January 2018 to confirm our findings.

**Results:** Thirty-five medical journals publishing original research from 13 publishers were included in the analysis. All 35 journals offered some form of open access with varying embargo periods of up to 12 months. Of these journals, 21 (60%) provided immediate open access with a CC BY licence under certain circumstances (e.g. to specific research funders). Of these 21, 20 only offered a CC BY licence to authors funded by non-commercial organizations and one offered this option to funders who required it.

**Conclusions:** Most leading medical journals do not offer to authors reporting commercially funded research an open access licence that allows unrestricted sharing and adaptation of the published material. The journals’ policies are therefore not aligned with open access declarations and guidelines. Commercial research funders lag behind academic funders in the development of mandatory open access policies, and it is time for them to work with publishers to advance the dissemination of the research they fund.

**Strengths and limitations of this study:**

- This manuscript includes a systematic analysis of open access policies of journals with a high impact factor, including society-owned journals, from multiple publishers.
- The open access policies of all journals analysed were clarified, and confirmation of our findings was received by email from 97% of the contacted journals.
- Open access policies of the journals and publishers analysed are subject to change, so the information presented here may not be current.
- By selecting journals with a high impact factor, our analysis does not include prestigious journals from specialized therapy areas and regional or non-English language journals, which may have lower impact factors.
- Although our study covers only a small number of journals, extending such a manual analysis to a greater number of journals without loss of detail and verification of all results would be cumbersome and inefficient by relying on traditional analysis tools.

## INTRODUCTION

Hundreds of billions of US dollars are invested in medical research by governments, charities and commercial organizations each year, with the aim of extending and improving human lives.^1^ Publication plays an important role in the dissemination of scientific innovation.^2 3^ However, translation of medical research into clinical practice is slow; one study has suggested that it takes an average of 17 years for research evidence to reach 50% adoption in clinical practice, with the longest delays occurring after successful publication of clinical trial results.^2 3^

Open access publishing has the potential to improve innovation and speed up its adoption. Complete access to research literature encourages viewing of more articles than partial access,^4 5^ and open access articles appear to be downloaded more often and receive more citations than subscription articles, indicating a greater academic impact.^6-9^ There is also evidence suggesting that open access articles have a broader societal impact based on Altmetric data that measure the amount of attention publications receive in the news media and social communication channels.^9-11^ Depending on the restrictiveness of its licensing, open access can facilitate public and commercial reuse of research results, which is beneficial for collaboration, education and innovation.^9^ Furthermore, enabling access to the full text of research articles increases transparency, which benefits the public by helping both doctors and patients to find complete and current evidence to inform treatment decisions, and by preventing potentially harmful decisions being made based on the abstracts of paywalled articles.^9 12-14^ It is worth noting that the publishing model used by a journal (i.e. open access or subscription) has no impact on the quality of articles published.^15 16^

‘Open access’ is a broad term that encompasses a range of definitions, from ‘free-to-read’ (full text available to read on demand, without charge to the reader) to ‘free-to-read and reuse’ (with the additional ability to reuse text, tables and figures in different formats). When a journal offers open access, it has wide leeway in the choice of policy or policies it will apply, using one of the Creative Commons licences that allow reuse under specific terms, or offering free-to-read access without a licence.^17^

The Open Access Scholarly Publishers Association and the Budapest Open Access Initiative recommend the Creative Commons Attribution (CC BY) licence because it allows sharing and adaptation of published materials for any purposes (both commercial and non-commercial), subject only to attribution of the original source.^17-19^ Common alternatives to the CC BY licence include CC BY Non-Commercial (CC BY-NC), which restricts commercial reuse; CC BY No Derivatives (CC BY-ND), which restricts adaptation; and CC BY-NC-ND, which restricts both (table S1).^17 20^

Open access with a Creative Commons licence is typically facilitated by article processing charges. Following payment of such a charge by the research author, institution or funder, open access articles with a Creative Commons licence are usually made available on the journal’s website at the time of publication in the publisher’s typeset format (Version of Record). Open access articles that do not include a Creative Commons licence at the time of publication typically involve an embargo period before the published articles are freely accessible and may allow access only to the accepted manuscript (a version that has not been edited and typeset by the journal), which is made available on the author’s institutional website or on PubMed Central/Europe PubMed Central without a requirement for payment.

There has been an increasing trend towards open access publishing over the last 20 years, and almost 50% of articles were published open access in 2015.^8^ Many academic and not-for-profit research funders now require the research they fund to be published open access.^9 21-26^ Prominently, the Wellcome Trust and the Bill & Melinda Gates Foundation insist on publishing with a CC BY licence to allow the broadest possible use.^21 23^ Commercial research funders, which fund approximately half of all medical research,^1 27 28^ have been more hesitant to require open access publishing but now commonly pay for open access when the option is available.^24^ In January 2018, Shire became the first commercial research funder to require all research manuscripts it funds to be published open access.^29 30^ At present, no commercial funder requires open access publishing with a CC BY licence.

We set out to clarify the open access variants provided by leading medical journals for research, in general and industry-funded research in particular, and establish the availability of the CC BY licence for industry-funded research.

## METHODS

Using Journal Selector (Sylogent, Newtown, PA, USA), we identified medical journals with a 2015 impact factor of at least 15.0 (accurate on 24 May 2017). To focus on journals publishing original medical research, we excluded journals that only publish review articles. We collected information on the open access variants provided by the included journals from their websites and by email contact when information was missing or unclear, making up to three attempts between 29 June 2017 and 26 July 2017.

For each journal, we recorded the following information:

- for immediate open access, whether a CC BY licence or other Creative Commons licence was provided
- for delayed open access, the length of embargo period for open access
- for both immediate and delayed open access, which version of the article would be available (published Version of Record or accepted).

For journals that provided a CC BY licence, we additionally collected information on:

- the requirements for obtaining a CC BY licence (e.g. dependence on funding source)
- article processing charges.

Between 6 December 2017 and 2 January 2018, we emailed the journals’ editorial offices requesting confirmation of our findings (table S2). Once open access variants were recorded, we categorized the most open variant provided by each included journal using our own classification, as shown in table 1.

**Table 1.** Categorization of journals based on the most open variant of open access offered

| Category | Version of article available | Embargo period | CC BY licence offered by the journal? |
| --- | --- | --- | --- |
| 1 | Published | None | Yes |
| 2 | Published | None | No |
| 3 | Published/accepted | ≤ 12 months | No |

CC BY, Creative Commons Attribution.

## RESULTS

### Included journals

Fifty-three journals listed in the Journal Selector database had a 2015 impact factor of at least 15.0 (figure 1). After 16 review journals and two non-medical journals were excluded, 35 journals from 13 publishers were included in this analysis. Of the 15 journals that were contacted to clarify information that was missing or unclear, 14 replied with clarification. Once all information was collected and tabulated, we received confirmation of our findings from 34 (97%) of the 35 journals.

**Figure 1.**
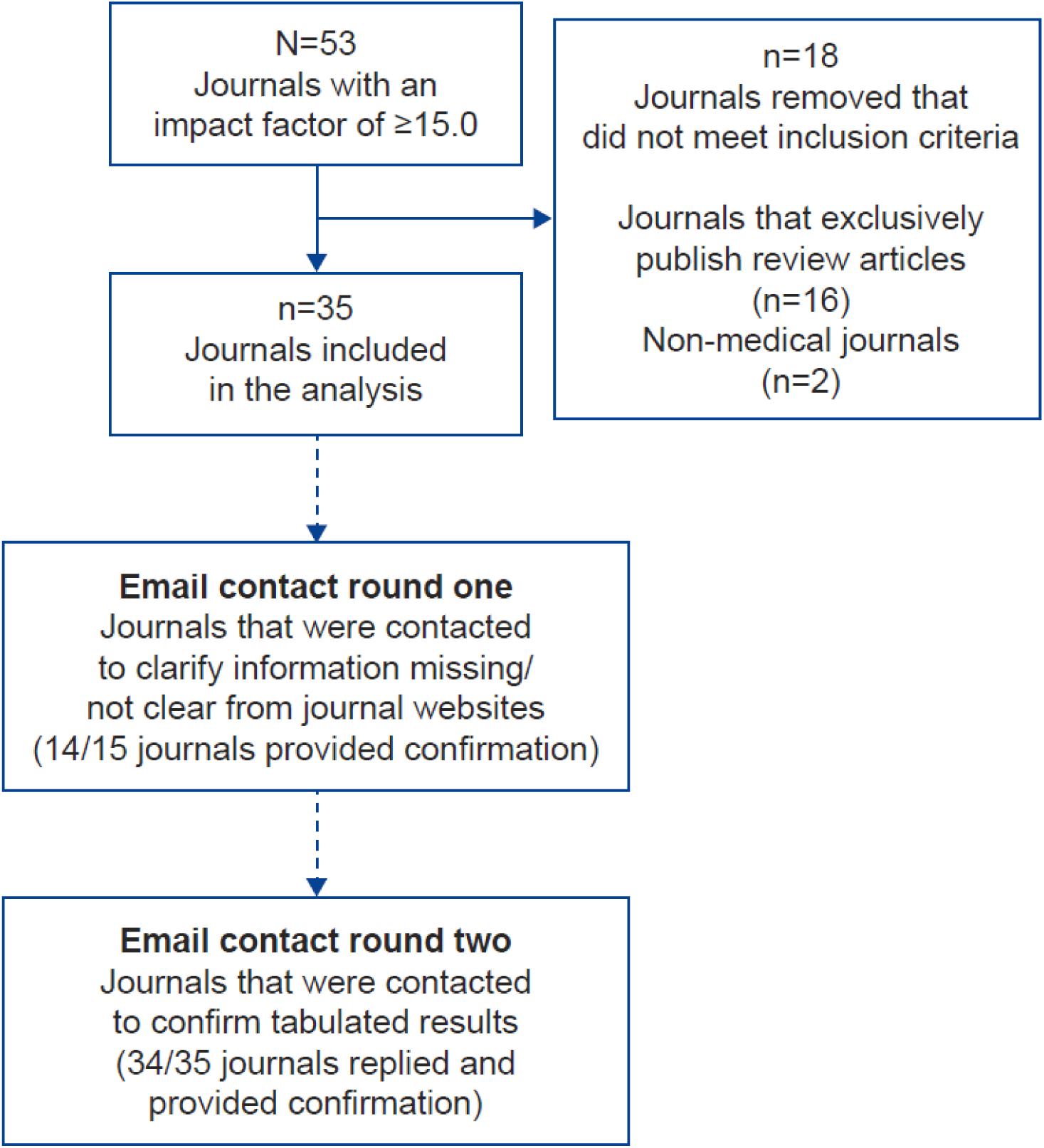
Flow chart of journals included in this study.

### Open access variants offered

Proportions of journals in each category of the most open variant of open access are shown in figure 2A. Immediate open access with a Creative Commons licence was provided by 21 (60%) of the 35 journals analysed. The types of Creative Commons licence available from these 21 journals under different circumstances were: CC BY from 21 journals (100%); CC BY-NC from 4 journals (19% of all journals offering CC BY); and CC BY-NC-ND from 18 journals (86% of all journals offering CC BY).

**Figure 2.**
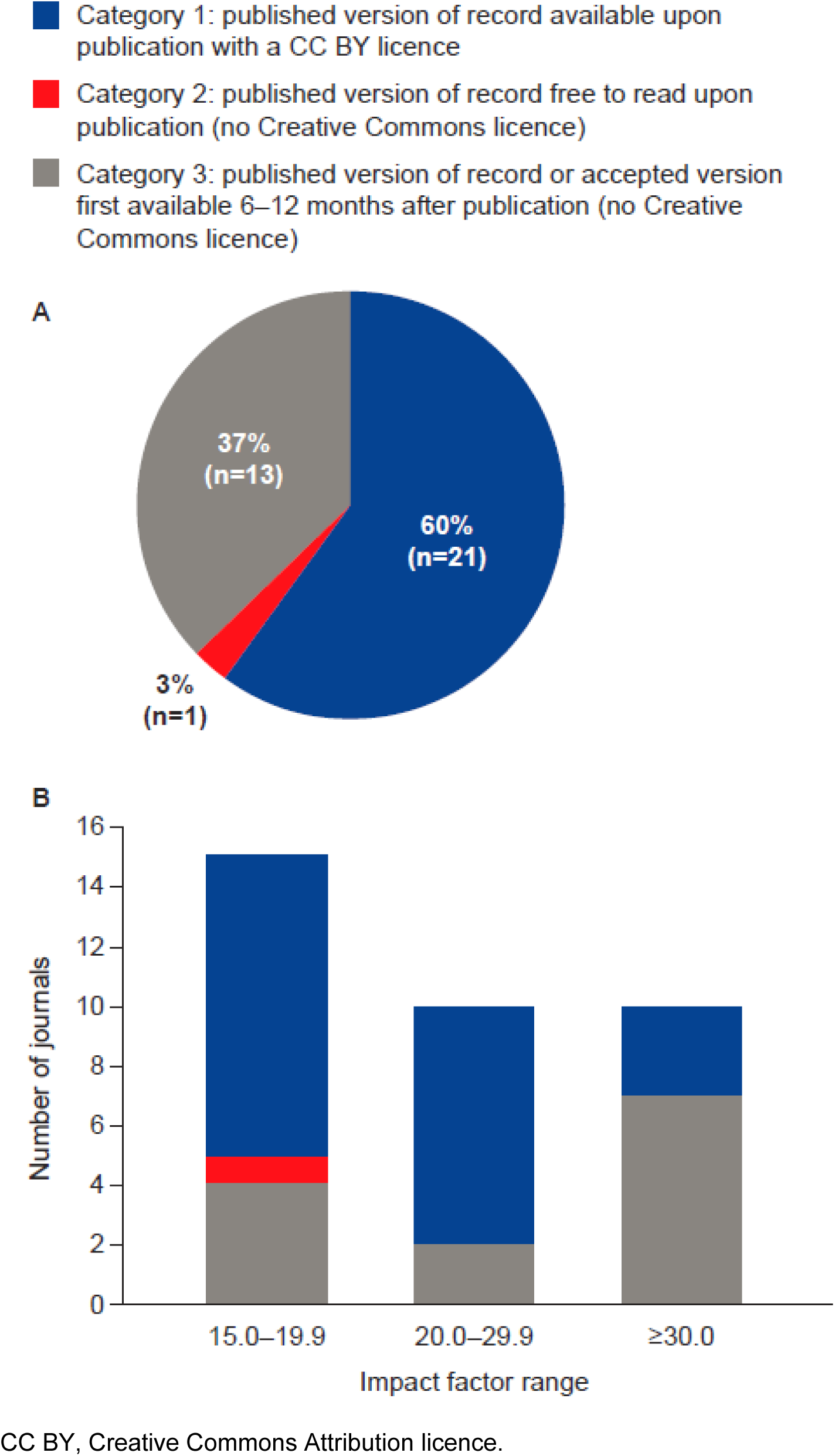
Medical journals categorized by impact factor and their most open variant of open access available (n=35). **(A)** Impact factor, ≥15.0; **(B)** Impact factors, 15.0– 19.9, 20.0–29.9 and ≥30.0.

When the 35 analysed journals were categorized by impact factor, immediate open access with a CC BY or other Creative Commons licence was provided by 10 (66%) of the 15 journals with an impact factor between 15.0 and 19.9, and 3 (30%) of the 10 journals with an impact factor over 30.0 (figure 2B).

All 14 journals, from six publishers, that did not provide open access with a Creative Commons licence provided access to different versions of the article either immediately, after a 6-month embargo period or after a 12-month embargo period under different circumstances (table 2).

**Table 2.**
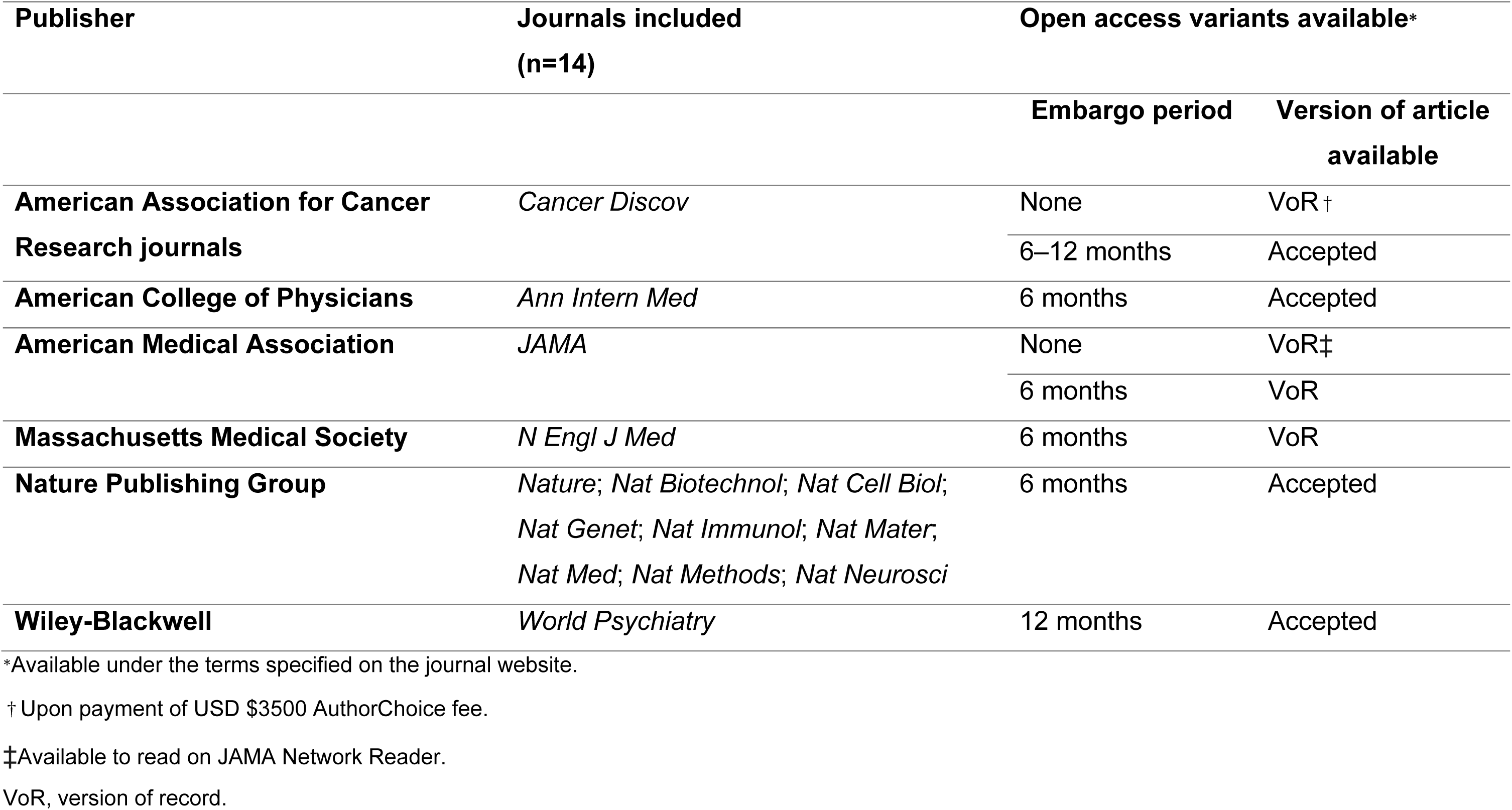
Access policies of journals with high impact factors that do not provide open access with Creative Commons licences.

| Publisher | Journals included<br>(n=14) | Open access variants available* |  |
| --- | --- | --- | --- |
|  |  | Embargo period | Version of article available |
| American Association for Cancer Research journals | *Cancer Discov* | None | VoR† |
|  |  | 6–12 months | Accepted |
| American College of Physicians | *Ann Intern Med* | 6 months | Accepted |
| American Medical Association | *JAMA* | None | VoR‡ |
|  |  | 6 months | VoR |
| Massachusetts Medical Society | *N Engl J Med* | 6 months | VoR |
| Nature Publishing Group | *Nature; Nat Biotechnol; Nat Cell Biol; Nat Genet; Nat Immunol; Nat Mater; Nat Med; Nat Methods; Nat Neurosci* | 6 months | Accepted |
| Wiley-Blackwell | *World Psychiatry* | 12 months | Accepted |

\*Available under the terms specified on the journal website.
† Upon payment of USD \$3500 AuthorChoice fee.
‡ Available to read on JAMA Network Reader.
VoR, version of record.

### The cost of open access with a CC BY licence

Of the 21 journals that offered a CC BY licence, 19 (90%) disclosed article processing charges on their websites. Across these journals, charges ranged from USD $3000 to $5000; the most common article processing charge was $5000 (in 13 [62%] of journals; figure 3). Details of the fees charged by the remaining two journals (10%) were not available from their websites because the details were only provided when the article was accepted.

**Figure 3.**
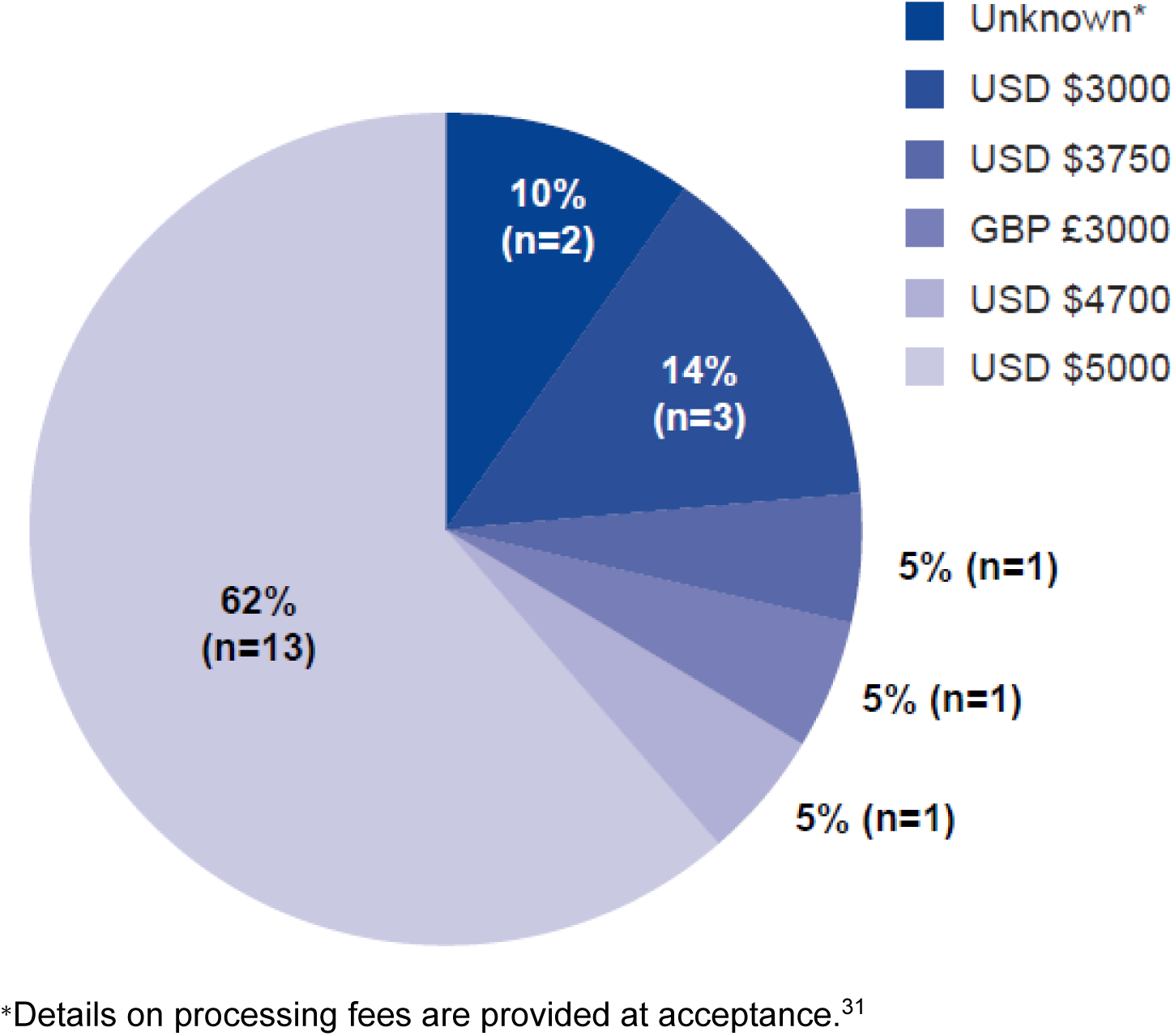
Article processing charges of journals that offer immediate open access with a CC BY licence (n=21).

### Relationship between funding source and the availability of open access variants

Table 3 shows the open access policies of the journals offering open access with a CC BY licence. Of the 21 journals listed, 20 journals allowed open access with a CC BY licence for research funded by specific non-commercial organizations, and only *The BMJ* offered it to organizations who required it, regardless of the nature of the funding source.

**Table 3.** Open access policies of journals with high impact factors that offer immediate open access with the CC BY licence (n=21) Creative Commons licences are shaded.

| Publisher | Journals included (n=21) | Open access variants available* |  |  | Funding requirements for obtaining open access with a CC BY licence |
| --- | --- | --- | --- | --- | --- |
|  |  | Embargo period | Creative Commons licence | Version of article available |  |
| American Association for the Advancement of Science | *Science*;<br>*Sci Transl Med* | None | CC BY | VoR | The American Association for the Advancement of Science “will allow authors funded by the Bill & Melinda Gates Foundation to publish their research with a CC BY licence”<sup>†</sup> |
|  |  | None | None | Accepted |  |
|  |  | 6 months | None | Accepted |  |
|  |  | 12 months | None | VoR |  |
| American Society of Clinical Oncology | *J Clin Oncol* | None | CC BY<br>CC BY-NC-ND | VoR | Creative Commons licences available only if funders are “academic institutions, not-for-profit organizations, philanthropic foundations or government agencies” |
|  |  | 6 months | None | VoR |  |
|  |  | 12 months | None | VoR |  |
| BMJ Publishing Group | *BMJ* | None | CC BY<br>CC BY-NC | VoR | CC BY licence available for authors “where the funder requires it” |
| Cell Press |  | None | CC BY | VoR |  |
|  | *Cancer Cell; Cell;* |  | CC BY-NC-ND |  | Creative Commons licences “available only to authors covered by a funding body agreement” (these non-commercial funding bodies are listed on the journal websites) |
|  | *Cell Metab;* | 12 months | None | Accepted |  |
|  | *Cell Stem Cell;* |  |  |  |  |
|  | *Immunity* |  |  |  |  |
| **Elsevier** | *Eur Urol;* | None | CC BY | VoR | Creative Commons licences are available to authors funded by specific funding bodies (these non-commercial funding bodies are listed on the journal websites) |
|  | *Gastroenterology;* |  | CC BY-NC-ND |  |  |
|  | *J Am Coll* | 6 months | None | VoR | Elsevier has established agreements and developed policies to allow authors who publish in Elsevier journals to comply with manuscript archiving requirements of various funding bodies (these non-commercial funding bodies are listed on the journal websites) |
|  | *Cardiol; Lancet;* |  |  |  |  |
|  | *Lancet Diabetes* |  |  |  |  |
|  | *Endocrinol;* |  |  |  |  |
|  | *Lancet Infect Dis;* |  |  |  |  |
|  | *Lancet Oncol;* |  |  |  |  |
|  | *Lancet Neurol;* |  |  |  |  |
|  | *Lancet Respir* |  |  |  |  |
|  | *Med* |  |  |  |  |
| **European Society of Cardiology** | *Eur Heart J* | None | CC BY | VoR | “RCUK/Wellcome Trust-funded authors...can use the CC BY licence for their articles” |
|  |  |  | CC BY-NC |  |  |
|  |  |  | CC BY-NC-ND |  |  |
|  |  | None | None | Accepted |  |
|  |  | 12 months | None | Accepted |  |
| **Lippincott**<br>**Williams &**<br>**Wilkins** | *Circulation* | None | CC BY<br>CC BY-NC<br>CC BY-NC-ND | VoR | “Note that authors funded by RCUK or the Wellcome Trust may choose the CC BY licence if they agree to pay the article processing charge and commercial reuse of the article is not a factor” |
|  |  | 6–12 months | None | Accepted |  |
| **Wiley-**<br>**Blackwell** | *CA Cancer J Clin* | None | CC BY<br>CC BY-NC<br>CC BY-NC-ND | VoR | “All RCUK and Wellcome Trust-funded authors will be directed to the CC BY licence” |
|  |  | 12–24 months | None | Accepted |  |

\* Available under the terms specified on the journal website.
†The American Association for the Advancement of Science's pilot open access partnership with the Gates Foundation concluded on 30 June 2018.<sup>31</sup>
CC BY, Creative Commons Attribution; NC, Non-Commercial; ND, No Derivatives; RCUK, Research Councils UK; VoR, version of record.

## DISCUSSION

Here, we present a systematic analysis of open access policies of journals with a high impact factor, including society-owned journals, from multiple publishers. We met our objective to clarify the open access policies of all journals analysed and received confirmation of our findings by email from 97% of the contacted journals. We found that all leading medical journals in this study provided some form of open access, but there was little consistency across their policies. Over half of the included journals provided a CC BY licence; however, with the exception of one journal, this option was only available only to authors funded by non-commercial organizations. One journal (*The BMJ*) allowed authors to obtain a CC BY licence when the work was supported by funders who required its use. Therefore, if pharmaceutical companies had a policy that required open access with a CC BY licence, the *The BMJ* would be suitable, and other journals might be inclined to change their policy.

Limitations of this study are that we investigated journals listed in the Journal Selector database with an impact factor of at least 15.0, and that, because impact factors and the open access policies of journals and publishers are subject to change, the information may not be current. Furthermore, by selecting journals with a high impact factor, our analysis does not include prestigious journals from specialized therapy areas and regional or non-English language journals, which may have lower impact factors. Although our study covers only a small number of journals, extending such a manual analysis to a greater number of journals without loss of detail and verification of all results would be cumbersome and inefficient by relying on traditional analysis tools. If more extensive mining of journal (meta)data becomes feasible, however, this study could be repeated for a bigger cohort of journals.

To our knowledge, this is the first report showing that the availability of open access options depends on the source of funding. Limitations on the availability of the CC BY licence depending on the research funder are not in line with statements such as the Budapest Declaration,^18^ the Berlin Declaration^32^ and the Bethesda Statement,^33^ which aim to provide end users with immediate access to research articles and to give them the opportunity to reuse material without restrictions. Furthermore, placing restrictions on access to medical research owing to its source of funding is not in line with the key principles of human research ethics laid out in the Declaration of Helsinki.^34^

Good Publication Practice 3 (GPP3) guidelines state that authors should take responsibility for the way research findings are published.^35^ In line with these recommendations, pharmaceutical companies can and, we believe, should advise authors to reach a consensus on which journal to publish with, to avoid predatory journals, and to adhere to sponsor guidelines and regulations. In the authors’ experience, some pharmaceutical companies already have internal guidelines recommending open access publishing, and one (Shire) now requires it.

Our research shows that one-third of the journals with a high impact factor do not offer immediate access to the published version of a manuscript upon publication, even though the open access policies of many funders with respect to embargo periods echo the recommendations set out by open access declarations worldwide.^18 21-23 26 32 33 36^ Of note, Horizon 2020, which is supported by the European Research Council, requires its beneficiaries to make publications open access no later than 6 months after the official publication date and to make every effort to allow for maximum reuse of the materials, whether that be copying, distributing, searching, linking, crawling, mining or some other use.^37 38^ Furthermore, cOAlition S, a group of national research funders with the support of the European Commission and the European Council, has committed to Plan S, the key principle of which is that scientific publications on research funded by participating national and European funders must be published open access by 2020.^36^ Under the terms of Plan S, authors must retain copyright of their publication with no restrictions, and all publications must be published under an immediate open licence (preferably CC BY) that fulfils the requirements defined by the Berlin Declaration.^36 37^

Policies vary between publishers but also across journals at the same publisher, and this is also the case for journals not included in this analysis, as shown, for example, by Taylor & Francis in their table of the policies of all their journals.^39^ Differences in policy have many underlying factors, including the choices of the journals’ academic editorial boards and societies. A potential disincentive to publishers offering CC BY licences to the pharmaceutical industry is the revenue generated from copyright fees and reprints. Permission to reproduce copyrighted materials can cost hundreds or even thousands of dollars; for example, the permission fee requested for reuse of a single table containing 40 words in the journal *American Family Physician* was $4400.^40^ Reprints can cost significantly more than permissions charges; for example, reprint sales from a single clinical trial can total $1 million or more, with a large profit margin.^41^

Research by Lundh *et al.*^42^ aimed to quantify reprint revenues as a proportion of journal income. Of the six journals investigated, the two European journals, *The BMJ* and *The Lancet*, disclosed the information requested. The editors of the US journals *Archives of Internal Medicine, Annals of Internal Medicine, JAMA* and the *New England Journal of Medicine* did not provide the data. For *The BMJ*, reprint revenues constituted 3% of its overall income; *The Lancet* obtained 41% of its revenue from reprints.^42^ In *The Lancet*, industry-funded publications constituted a large proportion of highly reprinted articles (63/88) compared with a sample of control articles from the same journal (23/88).^43^ The generation of revenue for publishers from the selling of reprints leaves publishers open to the criticism that bias can be introduced into editorial decisions.^42^ This concern could be addressed by a transition to open access publishing exclusively with a CC BY licence. However, such a transition may need to be managed.

Two of the journals included in our analysis, *Science* and *Science Translational Medicine*, both published by the American Association for the Advancement of Science, do not disclose article processing charges on their websites;^31^ instead, they provide this information upon their acceptance of an article. This practice does not comply with the Directory of Open Access Journals guidelines,^44^ which state that processing fees must be stated clearly on journal websites in a place that is easy to find for potential authors prior to submitting their manuscript. The practice is also common among predatory journals, potentially reinforcing perceptions held by some academics of the association between open access and predatory publishing.

We found that the open access policies of some journals precluded commercially funded research from being published open access, even after an embargo period and without a Creative Commons licence. Further analyses could therefore be undertaken to clarify the proportion of journals with this policy and the rationale behind this position. Future research could also focus on a larger cohort of journals than the current study, or on journals from a specific therapy area, to further clarify the use of open access variants in the medical publications landscape.

## CONCLUSIONS

The CC BY licence is recommended by open access declarations and funders of research as the optimal open access licence. Our analysis shows that although journals with a high impact factor provide some form of open access, they restrict commercially funded research from being published with the CC BY licence. Approximately half of all medical research is funded by the pharmaceutical industry,^1 27 28^ meaning that the research output cannot be reused or built upon if it is published in journals with a high impact factor without payment of additional fees, hampering research innovation and collaboration. However, there are concerns that a rapid transition to publishing exclusively with a CC BY licence will be difficult, given current processes and business models in scientific publishing.

The idea that open access to research articles is beneficial to all stakeholders in medical research and publishing is compelling. Open access publishing facilitates faster and more thorough disclosure of research, removes barriers for groups conducting systematic reviews, increases both the citation counts and Altmetric scores of publications, and benefits patient health by improving informed decision-making by doctors and patients.^9^ Commercial research funders lag behind non-commercial funders in the implementation of open access policies, and it is time for them to close the gap. Pharmaceutical companies should make clear their open access requirements, for example in a unified position statement, ideally aligned with open access declarations,^18 32 33^ the Horizon 2020 programme and Plan S,^36-38^ and the International Committee of Medical Journal Editors^45^ and GPP3^35^ guidelines, and then work together with publishers to realise the ultimate goal of improved access to medical research for all.

## Supporting information

## Acknowledgements

Robert Kiley (https://orcid.org/0000-0003-4733-2558) is Head of Open Research at the Wellcome Trust, London, UK, and contributed to the review of this manuscript. Paul Farrow (https://orcid.org/0000-0002-0569-9688) is an employee of Oxford PharmaGenesis, Oxford, UK, and contributed significantly to the review of this manuscript. Sarah Stokes (https://orcid.org/0000-0002-8761-8588) and Velissaria Vanna are employees of Oxford PharmaGenesis, Oxford, UK, and contributed to the review and editing of this manuscript. This work was presented as a poster at both the European Meeting of the International Society for Medical Publication Professionals (ISMPP) on 23 January 2018 and the Annual Meeting of ISMPP on 2 May 2018, and was posted to bioRxiv as a preprint on 22 January 2018 (https://www.biorxiv.org/content/early/2018/01/22/250613).

## Author contributions

Conceptualization, project administration, TE (https://orcid.org/0000-0003-0307-725X), TK (https://orcid.org/0000-0001-6152-7365), LS (https://orcid.org/0000-0001-6117-781X), AW (https://orcid.org/0000-0002-9354-6402); methodology, resources, investigation, formal analysis, TE; writing – original draft, TE and LS; visualization, TE; writing – review and editing, TE, TK, LS, AW, CW (https://orcid.org/0000-0003-3267-3990); supervision, TK, LS

