## Supplementary material for "Open access policies of leading medical journals: a cross-sectional study"

### Supplemental information

**Table S1** Definitions of CC licences commonly used by medical journals[17]

| Type | Definition |
|---|---|
| **CC BY** | Free to distribute and adapt the original work, even commercially, if the original creation and authors are credited |
| **CC BY-NC** | Free to adapt the original work non-commercially and, although derivative works must also acknowledge the authors and be non-commercial, they do not have to be licensed on the same terms |
| **CC BY-NC-ND** | Free to download the original work and share it if the authors are credited, but the work cannot be adapted or used commercially |

CC, Creative Commons; CC BY, Creative Commons Attribution; NC, Non-Commercial; ND, No Derivatives.
