## Supplementary material for "Open access policies of leading medical journals: a cross-sectional study"

### Supplemental information

**Table S2** Emails to and from journals

| Publisher | Journal (n=37) | Email question | Email response |
|---|---|---|---|
| **American Association for Cancer Research Journals** | *Cancer Discov* | (To:)<br>(Sent: 6 December 2017)<br><br>I have been looking on the AACR website for information about open access options for Cancer Discovery. Please could you confirm that the options available for this journal are:<br><br>Immediate open access: open access to the published article without a Creative Commons (CC) licence<br>Access 6–12 months after publication: green open access with no CC licence (accepted version uploaded to a repository – only if funded by a specific non-commercial organization) | (From: …@aacr.org)<br>(Received: 14 December 2017)<br><br>I have copied our archiving and open access policies below. You can also find these policies on our website (http://aacrjournals.org/content/authors/copyright-permissions-and-access). If you have any other questions, please let me know. |
| **American Association for the Advancement of Science (AAAS)** | *Science;*<br>*Sci Transl Med* | (To: mailto:)<br>(Sent: 6 December 2017)<br><br>I have been looking on the AAAS website for information about open access options for the *Science* and *Sci Transl Med* journals for research I am conducting. Please could you confirm that the options available for these journals are:<br><br>**Immediate open access:** CC BY licence (only if authors are funded by the Gates Foundation); green open access with no CC licence (accepted version uploaded to personal/institutional website)<br>**Access 6 months after publication:** green open access with no CC licence (accepted version uploaded to a repository)<br>**Access 12 months after publication:** green open | (From:)<br>(Received: 7 December 2017)<br><br>Thank you very much for getting in touch. This is essentially correct but I'm re-writing it a bit to clarify some points (I hope!).<br><br>For Science and Science Translational Medicine we offer a<br><br>Science Journals Default license. This is not a Creative Commons license. It is Green open access which means after publication the author may:<br><br>-post the accepted version of his paper on his personal or institution's repository<br>-after a 6 month embargo, the accepted version of the paper may be made available via your funding body archive or designated repository so long as the article copy is the restricted to non- |

| Publisher | Journal (n=37) | Email question | Email response |
|---|---|---|---|
| | | access with no CC licence (published version on journal website) | commercial research purposes<br>-after 12 months the paper is available for free from the off of journal website. This is the final published version.<br><br>If you are an author funded by the Gates foundation your paper will be released under the Creative Commons with attribution license (CC BY). Under a CC BY license the paper will be immediately available to read and download under the terms of the Creative Commons Attribution License (CC BY 4.0). Here is a link to the license: https://creativecommons.org/licenses/by/4.0/<br><br>I wanted to mention that our journal Science Advances currently only releases article under the CC licenses (CC BY and CC BY-NC). http://advances.sciencemag.org/content/article-licensing |
| **American College of Physicians** | *Ann Intern Med* | (To:)<br>(Sent: 6 December 2017)<br><br>I have been looking on the American College of Physicians website for information about open access options for Ann Intern Med. Please could you confirm that the options available for this journal are:<br><br>Immediate open access: open access with a Creative Commons (CC) licence is not available for this journal<br>Access 6 months after publication: green open access with no CC licence (accepted version uploaded to a repository – only if funded by NIH) | (From:)<br>(Sent 6 December 2017)<br><br>Yes this is correct. |
| **American Chemical Society** | *Acc Chem Res* | **Email 1:**<br>(To: ACS Publications Support [via website])<br>(Sent: 6 December 2017)<br><br>I have been looking on the ACS Publications website for information about open access options for the Acc Chem Res journal for research I am conducting. Please could you confirm that the options available for this journal are:Immediate open access: CC BY licence (any author can obtain this licence regardless of funding source); CC BY-NC-ND; open access with no | **Email 1:**<br>(From:)<br>(Sent: 6 December 2017)<br><br>Please note that all ACS Author Choice options you mentioned are available for the Accounts of Chemical Research.<br><br>http://pubs.acs.org/page/4authors/openaccess/index.html#achre4<br><br>For more information, please check the following links: |

| Publisher | Journal (n=37) | Email question | Email response |
|---|---|---|---|
| | | CC licence (published version)Access 12 months after publication: open access with no CC licence (published version)<br><br>**Email 2:**<br>(To:)<br>(Sent: 7 December 2017)<br><br>Thank you for your response and for providing extra information. Please could you clarify one point – is the Creative Commons CC BY licence only available to authors who are required to have one by their funding sponsors? Is the CC BY licence available to authors funded by commercial funders as well? | http://pubs.acs.org/page/4authors/authorchoice/options.html#optiona<br>http://pubs.acs.org/page/4authors/authorchoice/understanding_options.html#order<br>http://pubs.acs.org/pb-assets/documents/4authors/authorchoice_flowchart.pdf<br><br>**Email 2:**<br>(From)<br>(Sent: 8 December 2017)<br><br>Please note that Creative Commons CC BY licence is available to all authors, including the ones funded by commercial funders.<br><br>Some funders require a CC-BY license for all published research and that is the reason we point that information, but you may choose the CC-BY even if it is nor required by your funder.<br><br>Please check this decision tree and you will see that there is an option: "Do you need or would you like a Creative Commons license?"<br><br>http://pubs.acs.org/pb-assets/documents/4authors/authorchoice_flowchart.pdf |
| **American Medical Association** | *JAMA* | (Via: https://sites.jamanetwork.com/help/)<br>(Sent: 6 December 2017)<br><br>I have been looking on the JAMA website for information about open access options for JAMA. Please could you confirm that the options available for this journal are: Immediate open access: open access to the published article via the JAMA Network Reader (not subject to a Creative Commons [CC] licence) Access 6 months after publication: green open access with no CC licence (published version uploaded to journal website / repository). | (From …@jamanetwork.org)<br>(Sent: 6 December 2017)<br><br>Your query was forwarded to me. JAMA does not have an author-pay open access. As indicated in JAMA’s Instructions for Authors:<br><br>“All research articles are made free access online 6 months after publication on the journal website. All articles are made free access on The JAMA Network Reader on the day of publication.”<br><br>See https://jamanetwork.com/journals/jama/pages/instructions-for-authors#SecPublicAccess |

| Publisher | Journal (n=37) | Email question | Email response |
|---|---|---|---|
| **American Society of Clinical Oncology** | *J Clin Oncol* | (To:)<br>(Sent: 12 December 2017)<br><br>I have been looking on the American Society of Clinical Oncology website for information about open access options for the J Clin Oncol journal for research I am conducting. Please could you confirm that the options available for this journal are:<br><br>Immediate open access: CC BY and CC BY-NC-ND licences only available if funders are academic institutions, not-for-profit organizations, philanthropic foundations, or government agencies<br>Access 6 months after publication: green open access with no CC licence (published version uploaded to an institutional or funding body repository)<br>Access 12 months after publication: green open access with no CC licence (published version uploaded to a repository – for research funded by NIH and other US government agencies) | (From:)<br>(Sent: 13 December 2017)<br><br>I would say the following is correct:<br><br>Immediate open access: CC BY and CC BY-NC-ND licences only available if funders are academic institutions, not-for-profit organizations, philanthropic foundations, or government agencies<br>Access 6 months after publication: green open access with no CC licence (published version uploaded by the authors to an institutional or funding body repository)<br>Access 12 months after publication: standard JCO licence with federal funding (published version deposited by ASCO to PubMed Central (PMC) – for research funded by NIH and other US government agencies) |
| **BMJ Group** | *BMJ* | (To:)<br>(Sent: 13 December 2017)<br><br>I have been looking on the BMJ website for information about open access options for the BMJ journal for research I am conducting. Please could you confirm that the options available for this journal are:<br>Immediate open access to the published article: CC BY licence (only if authors are funded by non-commercial organizations that mandate open access publishing with this licence); CC BY-NC licence (default option) | (From: …@bmj.com)<br>(Sent: 14 December 2017)<br><br>Thanks for getting in touch. Those options you've suggested are pretty spot on - we offer CC BY-NC 4.0 as a default for all Research articles, although we offer a CC BY 4.0 licence on request when required by funders.<br>All other article types are not OA by default, but authors can request and pay for OA if required.<br>You can find out more about this here:<br>http://www.bmj.com/about-bmj/resources-authors/forms-policies-and-checklists/copyright-open-access-and-permission-reuse |
| **Cell Press** | *Cancer Cell;*<br>*Cell;*<br>*Cell Metab;*<br>*Cell Stem Cell;*<br>*Immunity* | (To:)<br>(Sent: 6 December 2017)<br><br>I have been looking on the Cell Press website for information about open access options for the Cancer Cell, Cell, Cell Metab, Cell Stem Cell and Immunity | (From: …@Cell.com)<br>(Sent: 15 December 2017)<br><br>What you have as a statement about open access for the titles you listed is correct. However only a subset of those funding bodies have a mandatory open access policy (Research Council UK, |

| Publisher | Journal (n=37) | Email question | Email response |
|---|---|---|---|
| | | journals. Please could you confirm that the options available for these journals are:<br><br>Immediate open access:CC BY and CC BY-NC-ND licences (only available to authors covered by a funding body agreement – listed below)<br>Access 12 months after publication:green open access with no CC licence (accepted version uploaded to a repository – for non-commercially funded research)<br><br>Funding body agreements for CC BY and CC BY-NC-ND licences:Arthritis Research UK, Arts and Humanities Research Council (UK), Biotechnology and Biological Sciences Research Council (UK), Bill and Melinda Gates Foundation, Bloodwise, British Heart Foundation (UK), Breast Cancer Now, Cancer Research UK, CAPES (Coordenação de Aperfeiçoamento de Pessoal de Nível Superior), Chief Scientist Office, Department of Defense (US), Department of Energy (US), Department of Health (UK), Dunhill Medical Trust, European Molecular Biology Laboratory (EMBL), European Research Council (ERC), Engineering and Physical Sciences Research Council (UK), Economic and Social Research Council (UK), Free University of Brussels (ULB), Gates Foundation, Howard Hughes Medical Institute (US), Hungarian Academy of Science, Joint Research Centre, Medical Research Council (UK), Motor Neurone Disease Association, Natural Environment Research Council (UK), National Institutes of Health (US), National Research Council Canada (NRC), Parkinson's UK, Research Councils (UK), Robert Koch Institute, Science and Technology Facilities Council (UK), Smithsonian Institution, Telethon (Italy), UK Department for International Development, United Nations University, VSNU (NL), Wellcome Trust (UK), World Bank, World Health Organization, Worldwide Cancer Research | Wellcome Trust, Gates Foundation) and those are the ones where authors can select open access. For example an NIH funded author is not offered a gold OA option but we do deposit the accepted article to PubMedCentral with a 12 month embargo. |

| Publisher | Journal (n=37) | Email question | Email response |
| --- | --- | --- | --- |
| **Elsevier** | *Lancet;*<br>*Lancet Diabetes Endocrinol;*<br>*Lancet Infect Dis;*<br>*Lancet Oncol;*<br>*Lancet Neurol;*<br>*Lancet Respir Med* | **Email 1:**<br>(To:)<br>(Sent: 6 December 2017)<br><br>I have been looking on individual journal websites for information about open access options for the Lancet, Lancet Diabetes Endocrinol, Lancet Infect Dis, Lancet Oncol, Lancet Neurol, and Lancet Respir Med journals. Please could you confirm that the options available for these journals are:<br><br>Immediate open access: open access to the published article with CC BY and CC BY-NC-ND licences (only available to authors funded by specific non-commercial organizations – listed below); green open access with optional attachment of CC BY-ND-ND licence (accepted version uploaded to personal/institutional websites or a repository)<br>Access 6 months after publication: green open access with no CC licence (published version uploaded to journal website and/or a repository)<br><br>Gold and green access are only offered to research funded by specific organizations: Arthritis Research UK, Austrian Science Fund, British Heart Foundation, Cancer Research UK, UK Chief Scientist Office, UK Department of Health UK, UK Department of International Development (DFID), Dunhill Medical Trust, Motor Neuron Disease Association, Parkinson's UK, one of the UK Research Councils, Telethon Italy, or Wellcome Trust; WHO (including International Agency for Research on Cancer [IARC]); Bill & Melinda Gates Foundation; Breast Cancer Now or Bloodwise; Worldwide Cancer Research<br><br>**Email 2:**<br>(To: …@elsevier.com)<br>(Sent: 7 December 2017) | **Email 1:**<br>(From: …@elsevier.com)<br>(Sent: 6 December 2017)<br><br>In the instructions for authors http://www.thelancet.com/authors/lancet/authorinfo there is a section dedicated to Open Access.<br><br>You can reference this for your research.<br><br>I hope this answers your question.<br>If there is anything else I can help you with, please let me know.<br><br>**Email 2:**<br>(From: …@elsevier.com)<br>(Sent: 7 December 2017)<br><br>Only papers submitted after a certain date and funded by specific organizations (see Open Access section of the Information for Authors for further details) can apply for Open Access.<br><br>I hope this helps you better. Let me know if you require further help. |

| Publisher | Journal (n=37) | Email question | Email response |
|---|---|---|---|
| | | Thank you for your response. Please could I get clarification on one point – is open access to the published article with the CC BY licence only available to authors funded by specific non-commercial organizations? Or is this licence available to all authors regardless of their funding source (e.g. authors funded by commercial organizations)? | |
| **Elsevier** | *Gastroenterology; J Am Coll Cardiol; Eur Urol* | **Email 1:**<br>(To:)<br>(Sent: 6 December 2017)<br><br>I have been looking on individual journal websites for information about open access options for the Gastroenterology, J Am Coll Cardiol, and Eur Urol journals. Please could you confirm that the options available for these journals are:<br><br>Immediate open access: open access to the published article with CC BY and CC BY-NC-ND licences (only available to authors funded by non-commercial organizations with established agreements with Elsevier); green open access with optional attachment of CC BY-ND-ND licence (accepted version uploaded to personal/institutional websites or a repository)<br>Access 6 months after publication: green open access with no CC licence (published version uploaded to journal website and/or a repository)<br><br>**Email 2:**<br>(To:)<br>(Sent: 7 December 2017)<br><br>Thank you for your response. Please could I ask for clarification on one point – is open access to the published article with the CC BY licence only available to authors funded by specific non-commercial organizations? Or is this licence available | **Email 1:**<br>(From:)<br>(Sent: 7 December 2017)<br><br>I can confirm that Open Access and Green Open Access for these journals are available. However, the embargo period for these is 12 months.<br><br>Should you require further assistance, please do not hesitate to contact me.<br><br>**Email 2:**<br>(From:)<br>(Sent: 8 December 2017)<br><br>I can confirm that, for these journals, CC BY license is only available to authors funded by specific non-commercial organizations. |

| Publisher | Journal (n=37) | Email question | Email response |
|---|---|---|---|
| | | to all authors regardless of their funding source (e.g. authors funded by commercial organizations)? | |
| **European Society of Cardiology** | *Eur Heart J* | **Email 1:**<br>(To:)<br>(Sent: 6 December 2017)<br><br>I have been looking on the European Society of Cardiology website for information about open access options for the European Heart Journal. Please could you confirm that the options available for this journal are:<br><br>Immediate open access: CC BY licence (only available for RCUK/Wellcome Trust-funded authors); CC BY-NC licence; CC BY-NC-ND licence; green open access with no CC licence (accepted version uploaded to personal/institutional websites, or submitted version uploaded anywhere provided there is a statement of acknowledgement of acceptance)<br>Access 12 months after publication: green open access with no CC licence (accepted version uploaded to a repository)<br><br>**Email 2:**<br>(To:)<br>(Sent: 13 December 2017)<br><br>*(Same message sent as before)*<br><br>**Email 3:**<br>(To: openaccess.oup.com)<br>(Sent: 2 January 2018) | **Email 1:**<br>(From: [automatic reply])<br>(Sent: 6 December 2017)<br><br>*(Automatic reply received detailing how to obtain journal permissions)*<br><br>**Email 2:**<br>**No response was received.**<br><br>**Email 3:**<br>(From:)<br>(Sent: 10 January 2018)<br><br>I hope you are well. Thank you for your email regarding the open access options available to authors publishing in European Heart Journal. Please be advised that authors publishing in European Heart Journal can chose to make their papers freely available via the gold open access method, by publishing their paper under an open access licence - the Creative Commons Attribution licence (CC-BY) for RCUK or Wellcome Trust funded authors, or the Creative Commons Attribution Non-Commercial licence (CC-BY-NC) for all other authors. Alternatively, authors can chose to make their papers freely available via the green open access method, by self-archiving a version of their paper. Authors may deposit the AoV (Author's Original Version) in personal/institutional websites, provided that once accepted they provide a statement of acknowledgment and, once published, this acknowledgment contains details of: volume number, issue number, DOI and a link to the published article. Or authors may |

| Publisher | Journal (n=37) | Email question | Email response |
|---|---|---|---|
| | | *(Same message as above was sent)* | deposit the AM (Accepted Version) on a personal webpage (excluding commercial websites) or institutional repository, on the proviso that it is not made publicly available until 12 months after publication. Further details as to how authors publishing in European Heart Journal can make their articles available via the green OA method can be found here: https://academic.oup.com/journals/pages/access_purchase/rights_and_permissions/self_archiving_policy_b. |
| **Lippincott Williams & Wilkins** | *Circulation* | **Email 1:**<br>(To:)<br>(Sent: 6 December 2017)<br><br>I have been looking on the Circulation website for information about open access options for the Circulation journal. Please could you confirm that the options available for this journal are:<br><br>Immediate open access: CC BY licence (only available to authors funded by RCUK or Wellcome Trust if commercial reuse isn't a factor); CC BY-NC licence; CC BY-NC-ND licence<br>Access 6–12 months after publication: green open access with no CC licence (publisher will deposit accepted version to a repository if funded by NIH, HHMI or The Wellcome Trust)<br><br>**Emails 2 and 3:**<br>*(Same message as above was sent to the same email address on 13 December 2017 and 2 January 2018)* | **Email 1:**<br>**No response was received.**<br><br>**Email 2:**<br>**No response was received.**<br><br>**Email 3:**<br>(From:)<br>(Sent: 2 January 2018)<br><br>The Editorial Office is closed until January 2nd. We will respond at our earliest convenience.<br>Happy Holidays and Happy New Year from the Circulation team. |
| **Massachusetts Medical Society** | *N Engl J Med* | (To:)<br>(Sent: 2 January 2018)<br><br>I have been looking on the journal website for information about open access options for N Engl J Med. Please could you confirm that the options available for this journal are:<br><br>Immediate open access: open access with a Creative | (From: …@mms.org)<br>(Sent: 4 January 2018)<br><br>Thank you for your email. In regards to your question, here is the NEJM information requested:<br><br>Six months after publication, NEJM makes full texts of all research articles available free of charge at NEJM.org. Certain materials may be released sooner at the discretion of NEJM |

| Publisher | Journal (n=37) | Email question | Email response |
|---|---|---|---|
| | | Commons (CC) licence is not available for this journal<br>Access 6 months after publication: green open access with no CC licence (published version uploaded to a repository) | editors. In particular, articles addressing matters of immediate interest to the public health are free from the time of publication. NEJM makes unrestricted immediate online access free to more than 90 low-income countries. NEJM does not offer an author-pays model, and there is no charge to submit or to publish in NEJM.<br>If research supporting an article was funded by the U.S. National Institutes of Health (NIH), the Wellcome Trust, or another not-for-profit organization that requires authors to submit to a publicly available, not-for-profit (non-institutional) repository, NEJM will submit PDFs of the published version to the NIH Manuscript Submission System or Europe PubMed Central to be released six (6) months after the article is published. |
| **Nature Publishing Group** | *Nature;*<br>*Nat Biotechnol;*<br>*Nat Cell Biol;*<br>*Nat Genet;*<br>*Nat Immunol;*<br>*Nat Mater;*<br>*Nat Med;*<br>*Nat Methods;*<br>*Nat Neurosci* | (To:)<br>(Sent: 6 December 2017)<br><br>I have been looking on Nature journals' websites for information about open access options for the following journals:<br><br>Nature<br>Nat Biotechnol<br>Nat Cell Biol<br>Nature Genetics<br>Nat Immunol<br>Nat Mater<br>Nat Med<br>Nat Methods<br>Nat Neurosci<br><br>Please could you confirm that the options available for these journals are:<br><br>Immediate open access: open access with a Creative Commons (CC) licence is not available for this journal<br>Access 6 months after publication: green open access with no CC licence (accepted version uploaded to a repository or shared by SharedIt content-sharing) | (From: …@springernature.com)<br>(Sent: 10 December 2017)<br><br>Thank you for contacting Nature Research about Open access options for Nature journals.<br><br>There are no paid immediate Open Access (OA) options available in this journal as Nature and the Nature Research journals support green OA deposition rather than gold OA publication, with the exception of Nature Communications, which is an open access title.<br><br>Open access through manuscript deposition<br><br>Authors publishing in Nature and in all other Nature Research journals are able to make the accepted version of their manuscript openly available six months after publication through self-archiving in the repository of their choice (see the full details of our self-archiving policy). Please note that version which may be archived is the manuscript accepted following peer-review and author revision, but prior to copyediting and typesetting by Nature Research.<br><br>We also offer a free Nature Research Manuscript Deposition Service of original research articles published in Nature, the |

| Publisher | Journal (n=37) | Email question | Email response |
|---|---|---|---|
| | | | Nature Research journals, and many of our society and academic journals. Corresponding authors whose funders have agreements with PubMed Central, Europe PubMed Central, and PubMed Central Canada may opt in to this service during submission. Nature Research will archive the accepted version of the manuscript on behalf of the author, and it will be made publicly accessible six months after publication with links back to the journal's website. For further information, please see here.<br><br>Journals with immediate OA option<br><br>If you are considering publishing an article via immediate (gold) OA in the future please do consult our list of journals with a paid open access option. Articles published in these journals can be made OA immediately on publication through payment of an article-processing charge (APC). Our funding support service can assist you in locating potential sources of APC funding. |
| **Wiley-Blackwell** | *Adv Mater* | (To:)<br>(Sent: 6 December 2017)<br><br>I have been looking on the journal website for information about open access options for the Advanced Materials journal. Please could you confirm that the options available for this journal are:<br><br>Immediate open access: CC BY licence (only available to authors with licencing mandates); CC BY-NC licence; CC BY-NC-ND licence; green open access with no CC licence (submitted version uploaded to a not-for-profit repository or to personal/institutional website)<br>Access 12 months after publication: green open access with no CC licence (accepted version uploaded to a not-for-profit repository or to personal/institutional website) | (From:)<br>(Sent: 11 December 2017)<br><br>Green open access is available in accordance with our copyright transfer agreement which is quoted at the end of this email.<br><br>Your information on paid open access is correct. In addition the accepted version of NIH-funded manuscripts is automatically indexed on Pubmed central after the 12-months embargo has expired. |
| | *CA Cancer J Clin* | **Email 1:**<br>(To:) | **Email 1:**<br>(From:) |

| Publisher | Journal (n=37) | Email question | Email response |
|---|---|---|---|
| | | (Sent: 6 December 2017)<br><br>I have been looking on the journal website for information about open access options for CA Cancer J Clin. Please could you confirm that the options available for this journal are:<br><br>Immediate open access: CC BY licence (only available to authors funded by RCUK or The Wellcome Trust); CC BY-NC licence; CC BY-NC-ND licence; green open access with no CC licence (published or submitted versions uploaded to a repository)<br>Access 12–24 months after publication: green open access with no CC licence (accepted version uploaded to a repository)<br><br>**Email 2:**<br>(To:)<br>(Sent: 2 January 2018)<br><br>I just wanted to follow up on my previous email. Please could I ask for clarification on the availability of the CC BY licence for CA Cancer J Clin? Is the CC BY licence only available to authors funded by specific non-commercial organizations? Or is this licence available to all authors regardless of their funding source (e.g. authors funded by commercial organizations)? | (Sent: 11 December 2017)<br><br>Thank you for your recent correspondence regarding the above journal title.<br><br>Please be advised Open Access are options for the above journal title, but as the journal is freely accessible, we have only ever had one article incorporate a CCBY license.<br><br>**Email 2:**<br>(From:)<br>(Sent: 3 January 2018)<br><br>Thank you for your follow up email regarding CC-BY license.<br><br>For this journal, CC-BY is only offered in compliant with your funder's policy. When signing your license agreement in Author Services, please select your mandated funder to ensure you are offered the appropriate license compliant with your funder's policy. Not indicating a funder with a license mandate during this process will lead you to sign either CC-BY-NC or CC-BY-NC-ND. |
| | *World Psychiatry* | **Email 1:**<br>(To:)<br>(Sent: 6 December 2017)<br><br>I have been looking on the journal website for information about open access options for World Psychiatry. Please could you confirm that the options available for this journal are:<br><br>Immediate open access: open access with a Creative | **Email 1:**<br>(From:)<br>(Sent: 7 December 2017)<br><br>Thank you for your recent correspondence regarding the above journal title.<br><br>Please be advised World Psychiatry is free to view and has been free to view since moving to Wiley in 2013 however is is not an Open Access title. |

| Publisher | Journal (n=37) | Email question | Email response |
|---|---|---|---|
| | | Commons (CC) licence is not available for this journal; green open access with no CC licence (submitted version uploaded to not-for-profit repository and personal/institutional website)<br>Access 12 months after publication: green open access with no CC licence (accepted version uploaded to not-for-profit repository and personal/institutional website)<br><br>**Email 2:**<br>(To:)<br>(Sent: 7 December 2017)<br><br>Thank you for your response. Please could I ask for clarification on one point – is it possible to purchase open access with a CC BY licence for individual articles with this journal? And if so, is the CC BY licence only available to authors funded by specific non-commercial organizations? Or is this licence available to all authors regardless of their funding source (e.g. authors funded by commercial organizations)? | **Email 2:**<br>(From:)<br>(Sent: 12 December 2017)<br><br>Thank you for your patience while we are checking into this.<br><br>Please be advised that there is no open access option available for this title as it is entirely free to read.<br><br>We hope this information helps. Please do not hesitate to contact us if you require any further assistance. |

Personal email addresses have been censored.
